# miR-486-3p ameliorates aging-related impaired angiogenesis in adult moyamoya disease by targeting Forkhead Box O4

**DOI:** 10.1101/2025.05.07.652777

**Authors:** Junda Chen, Yunyu Wen, Siyuan Chen, Tinghan Long, Fangzhou Chen, Zhibin Wang, Guozhong Zhang, Mingzhou Li, Shichao Zhang, Huibin Kang, Wenfeng Feng, Gang Wang

## Abstract

**Objective:** Investigate miR-486-3p’s role in alleviating age-related angiogenic decline in Moyamoya disease (MMD) by regulating senescent endothelial cells.

**Methods:** Clinical analysis of 151 MMD patients correlated age with postoperative angiogenesis (DSA grading). Senescent HUVECs (SA-β-gal>80%) exhibited elevated SASP factors (IL-6/IL-8/MCP-1). Functional assays (EdU/Transwell/Matrigel) and bioinformatics identified miR-486-3p targets, validated via luciferase/Western blot.

**Results:** Patients ≥35 years had 3.67-fold higher risk of poor angiogenesis (OR=3.67). Senescent HUVECs showed 10-16-fold higher SASP secretion (p<0.01). miR-486-3p overexpression increased proliferation (EdU+25%), migration (1.7-fold), and tube formation (+40% branches) in senescent cells (p<0.01) and enhanced angiogenesis in vivo (p<0.001). miR-486-3p directly targeted FOXO4, reducing its luciferase activity (−35%) and protein levels (−37%) (p<0.01), suppressing SASP.

**Conclusion:** miR-486-3p enhances post-revascularization angiogenesis in MMD by inhibiting FOXO4-mediated SASP, offering a therapeutic target and predictive biomarker.

## Introduction

Moyamoya disease is a cerebrovascular disorder characterized by chronic stenosis of the terminal portion of the internal carotid artery and the proximal segments of the circle of Willis, accompanied by the formation of an abnormal vascular network at the base of the brain[1],The pathogenesis of this disease remains unclear. The incidence rate in Asian populations is significantly higher than that in European and American populations. In China, the incidence has shown an increasing trend in recent years, with an annual rebleeding rate of 3.2%-16.4% among patients receiving conservative treatment.[2-7],Significantly increases both individual risk and societal burden. Current therapeutic approaches primarily focus on cerebral revascularization surgery.[8, 9],Indirect revascularization achieves compensatory blood supply through angiogenesis between the superficial temporal artery and middle meningeal artery, yet exhibits considerable variability in surgical efficacy, with postoperative adequate angiogenesis rates ranging from 42.2% to 84.8% in patients.[10-18],with adults demonstrating significantly lower rates compared to children, suggesting age-related factors influence angiogenesis efficacy.

Post-indirect revascularization, the intracranial-extracranial neovascular anastomosis process occurs within the cerebrospinal fluid (CSF) environment. Researchers have provided critical evidence indicating that exosomal miRNAs in CSF may reflect cerebral pathophysiological characteristics.[19] Building upon this foundation, our research team previously compared cerebrospinal fluid microRNA profiles between good and poor outcome groups following indirect revascularization. The results revealed significant differential expression of miR-486-3p, miR-25-3p, miR-92a-3p, and miR-155-5p between the groups, identifying these miRNAs as potential predictors of collateral circulation formation in Moyamoya disease patients post-indirect revascularization.[20],Consequently, our research focus has shifted to miR-486-3p, a previously uncharted microRNA in this context.

In this study, we demonstrate that miR-486-3p directly targets and suppresses FOXO4 expression, thereby reducing the secretion of senescence-associated secretory proteins (SASP) in senescent endothelial cells and restoring their angiogenic capacity. This work not only identifies miR-486-3p as a predictive biomarker for post-temporalis muscle attachment angiogenesis, but also elucidates the mechanistic role of the miR-486-3p/FOXO4 axis, providing potential therapeutic targets for enhancing vascular regeneration following surgical intervention.

## Materials and Methods

### 2.1 Ethical Statement

This study was approved by the Ethics Review Committee of Nanfang Hospital, Southern Medical University (Approval No.: NFEC-202302-K6-02), with written informed consent obtained from all participants. Experimental procedures strictly adhered to the guidelines of the Experimental Animal Ethics Committee at Southern Medical University. Mice were housed in specific pathogen-free (SPF) facilities with ad libitum access to food and water, undergoing 7-day acclimatization feeding prior to surgical procedures.

### 2.2 Patient Information Collection and Analysis

A single-center retrospective cohort study design was employed. The study enrolled 151 MMD patients (153 affected hemispheres) who underwent hospitalization in the Department of Neurosurgery at our institution from January 2014 to December 2022. Postoperative neovascularization was assessed using the Matsushima grading system: Grade A: Neovascular coverage ≥2/3 of the MCA territory (including frontoparietal lobe and temporo-occipital junction)

Grade B: Collateral compensation in 1/3-2/3 MCA territory (delimited by central sulcus)

Grade C: Compensatory range <1/3 or punctate neovascularization only

Grades A+B were categorized as “Good” outcomes, while Grade C denoted “Poor” outcomes.

### 2.3 Cell culture and senescence induction

Human umbilical vein endothelial cells (HUVECs) were purchased from the Shanghai Cell Bank of the Chinese Academy of Sciences. These cells were cultured in DMEM medium supplemented with 10% fetal bovine serum (FBS) at 37°C under 5% CO_2_ with saturated humidity. A replicative senescence model was established through continuous passaging. In this model, HUVECs exceeding 29 passages were classified as “senescent” cells, while those with fewer than 8 passages were defined as “normal” cells. Population doubling (PD) was calculated using the formula: PD = log_2_(F/I), where F represents the final cell count and I the initial cell count. Population doubling level (PDL) was determined by: PDLend = PDLinitial + PD.

### 2.4 Cell transfection

Sangon Biotech (Shanghai) synthesized hsa-miR-486-3p mimics, hsa-miR-486-3p inhibitors, negative control (NC), and Forkhead Box O4 (FOXO4)-specific siRNA. Overexpression plasmids and vector plasmids were provided by Umine-Bio. Co. Transfection of these oligonucleotides into HUVECs was performed using Lipofectamine 2000 reagent (Thermo Fisher Scientific) according to the manufacturer’s protocols.

### 2.5 Senescence-associated B-galactosidase (SA-β-gal) staining

SA-β-galactosidase staining was performed using a Senescence Detection Kit (Beyotime Biotechnology, Shanghai, China) following the manufacturer’s protocol. Briefly, cells were washed with PBS and fixed in SA-β-gal fixation solution at room temperature for 15 min. Subsequently, cells were rinsed three times with PBS and incubated with working solution at 37°C overnight. Images were captured under a Leica inverted microscope (Leica Microsystems GmbH) at 100× magnification. The percentage of positively stained cells was quantified by counting cells in six randomly selected microscopic fields.

### 2.6 BrdU incorporation assay

Cell proliferation was assessed using the 5-ethynyl-2’-deoxyuridine (EdU) incorporation assay. Briefly, cells were incubated with 10 μM EdU working solution in complete medium at 37°C for 2 hours, followed by fixation with 4% paraformaldehyde for 15 minutes. Fixed cells were permeabilized with 0.3% Triton X-100 in PBS and incubated at room temperature for 15 minutes. Click reaction was performed by adding Click Additive Solution under light-protected conditions for 30 minutes at room temperature. Nuclear counterstaining was conducted using Hoechst 33342. Fluorescent images were acquired with a Nikon inverted microscope (Nikon Instruments Inc.) at 100× magnification.

### 2.7 Transwell cell migration assays

HUVECs were resuspended in DMEM and quantified. A total of 10,000 cells in 200 μl serum-free DMEM were seeded into the upper chamber of a Transwell system. The lower chamber was loaded with 600 μl complete culture medium containing 10% FBS as chemoattractant. Following 12-hour incubation at 37°C, non-migratory cells in the upper chamber were removed using sterile cotton swabs. Migrated cells adherent to the membrane underside were fixed with 4% paraformaldehyde (PFA), stained with 0.1% crystal violet for 30 minutes, and quantified under an Olympus inverted microscope (Olympus Corporation).

### 2.8 In vitro tube formation assay

Matrigel (Corning Incorporated, USA) was thawed overnight at 4°C, with pre-cooled 96-well plates and pipette tips maintained on ice. Each well was coated with 50 μL Matrigel and polymerized at 37°C for 30 min (all experiments performed in triplicate). Transfected HUVECs were harvested 48h post-transfection, resuspended in complete medium (10% FBS), and quantified. A total of 15,000 cells were seeded per well in 100 μL complete medium (10% FBS), followed by 6-hour incubation at 37°C. Tubular structures were visualized using an Olympus IX73 inverted microscope (Olympus Corporation, Japan) equipped with a DP80 dual CCD camera. Quantitative analysis was performed using the Angiogenesis Analyzer plugin in ImageJ software (NIH, v1.53t).

### 2.9 Bioinformatics

We selected miR-486-3p’s target gene using TargetScan (www.targetscan.org/) and miRBase (www.mirdb.org/). Target genes were downloaded and generated into two Excel tables. The predicted target genes in the two tables were then analyzed by intersection analysis using the online Venn Diagramming site (http://bioinformatics.psb.ugent.be/webtools/Venn/). DAVID (https://david.ncifcrf.gov/) was used to analyze GO and KEGG signaling pathways using the intersection. After the analysis, all the data were saved, and the resulting table was searched for angiogenesis-related GO analysis and KEGG signaling pathway analysis, and the resulting related target genes were the possible target genes of miR-486-3p.

### 2.10 Western blotting analysis

Following cell lysis, protein concentration was quantified using RIPA buffer (Sigma-Aldrich, R0278). Samples were resolved via SDS-PAGE at 80 V for 2 hours, followed by electrophoretic transfer onto PVDF membranes (Millipore, IPVH00010; Billerica, MA, USA) at 320 mA for 100 minutes. Membranes were blocked with 5% bovine serum albumin (BSA; Solarbio Life Sciences, A8020) in Tris-buffered saline containing 1% Tween-20 (TBST; Sigma-Aldrich, P9416) for 1 hour. Primary antibodies were incubated with membranes at 4°C overnight. HRP-conjugated secondary antibodies (Cell Signaling Technology, Danvers, MA, USA) were applied at room temperature for 1 hour. Immunoreactive bands were visualized using Immobilon ECL Ultra Western HRP Substrate (Millipore, WBULS0500) on a Tanon-5500 Chemiluminescence Imaging System (Tanon Science & Technology, Shanghai, China).

#### Antibody Sources

Anti-CD18 (Catalog:10554-1-AP), Anti-FOXO4 (Catalog:21535-1-AP), Anti-GAB1 (Catalog:26200-1-AP), Anti-MEKK3 (Catalog:21072-1-AP), Anti-DLL1 (Catalog: 28544-1-AP), Anti-GAPDH (Catalog: 10494-1-AP): Wuhan San Ying Biotechnology (Wuhan, China)

### 2.11 Luciferase reporter assay

FOXO4-MT (mutant type) and FOXO4-WT (wild type) plasmids were procured from Wuhan GeneCreate Biotech Co., Ltd. (Wuhan, China). Plasmid transfection in 293T cells was performed under four experimental conditions:

FOXO4-MT alone

miR-486-3p mimics + FOXO4-WT

NC mimics + FOXO4-MT

miR-486-3p mimics + FOXO4-MT

Relative luciferase activity was quantified using the Dual-Luciferase Reporter Assay System (Promega, R41128) according to the manufacturer’s protocol.

### 2.12 In vivo Matrigel plug assay

A total of 300 μL Matrigel (Corning, Cat. No. 364263) was mixed with either miR-6760-5p agomir or antiagomir (Uming Biotechnology Co., Ltd., Guangzhou, China) and injected into the inguinal region of 8-week-old C57BL/6J mice. Following a 7-day incubation period, mice were euthanized under anesthesia, and Matrigel plugs were excised and fixed in 4% paraformaldehyde (PFA). Vascular protein content within the plugs was quantified using a Hemoglobin Assay Kit (Solarbio, Beijing, China) according to the manufacturer’s instructions.

### 2.13 Statistical analysis

Mean ± standard deviation (mean ± SD) from multiple independent experiments were reported, with differences among groups determined using one-way ANOVA in SPSS 24. 0. In addition, a Benjamini-Hochberg correction was applied to all p-values in order to control for false discovery rate (FDR).A significance level of p<0. 05 was considered significant.Using Graphpad Pism 9.5 for plotting

## Results

### 3.1 Age-Related Angiogenesis Outcomes in Moyamoya Disease

From clinical observations, pediatric patients demonstrated superior outcomes following indirect revascularization compared to adults, with reported success rates ranging from 42.2% to 84.8% in the literature (Supplementary Table 1). This study enrolled 151 Moyamoya disease (MMD) patients (78 females, 73 males; sex ratio 1.07:1), with a mean surgical age of 41.63 ± 13.45 years (range: 5–65 years), including 6.5% (n=10) adolescents under 18 years. Traditional vascular risk factors were notably less prevalent than in general cerebrovascular disease populations: hypertension (2.6%, n=4), smoking history (7.3%, n=11), and alcohol use (4.6%, n=7). Metabolic comorbidities included dyslipidemia (1.3%, n=2), diabetes (1.3%, n=2), and thyroid dysfunction (0.7%, n=1).(Table 1)

**Table 1.** Baseline Characteristics of 153 Patients.

| Characteristics | Value |
| --- | --- |
| Sex ratio (F/M) | 78:73 |
| Age at operation, mean ± SD, y | 41.63±13.45 |
| Age |  |
| <18 years | 10(6.5%) |
| History of risk factors |  |
| Hypertension | 4 (2.6%) |
| Smoking | 11 (7.3%) |
| Alcohol use | 7 (4.6%) |
| Thyroid disease | 1 (0.7%) |
| Hyperlipidemia | 2 (1.3%) |
| Diabetes | 2 (1.3%) |
| Surgical modalities |  |
| STA-MCA+EDMS | 110(71.9%) |
| STA-MCA+EDS | 36 (23.5%) |
| EDMS | 6 (4.0%) |
| EAS | 1 (0.7%) |
| DSA follow-up, mean±SD, mons | 9.73±7.02 |

The primary revascularization technique was superficial temporal artery-middle cerebral artery (STA-MCA) anastomosis combined with encephalo-duro-myo-synangiosis (EDMS) (71.9%, n=110), followed by STA-MCA with encephalo-duro-synangiosis (EDS) (23.5%, n=36). Pure EDMS (4.0%, n=6) and encephaloduro-arterio-synangiosis (EAS) (0.7%, n=1) were rarely utilized. Postoperative digital subtraction angiography (DSA) follow-up averaged 9.73 ± 7.02 months, with 55.0% (n=83) undergoing imaging assessments beyond 12 months. The cohort exhibited a bimodal age distribution, with a secondary peak in disease onset at 45–55 years (38.4%).

Using Matsushima grading, the good revascularization group (n=123) and poor group (n=30) differed significantly in baseline characteristics. The good group had a younger mean age (40.3 ± 13.9 vs. 47.0 ± 9.6 years; independent t-test, p = 0.014). Age-stratified analysis revealed that patients ≥35 years accounted for 90% (27/30) of the poor group versus 71.5% (88/123) in the good group (Fisher’s exact test, p = 0.036), identifying advanced age (≥35 years) as an independent risk factor for poor angiogenesis (OR = 3.67, 95% CI = 1.12–12.05). Sex distribution did not differ statistically (51.2% males in good vs. 36.6% in poor group; p = 0.16).(Table 2)

### 3.2 Cellular Senescence Impairs Endothelial Angiogenic Function via Autocrine-Paracrine Mechanisms

To establish a stable model of endothelial cell senescence, human umbilical vein endothelial cells (HUVECs) were induced into replicative senescence through serial passaging (up to passage 15). Senescence was confirmed via β-galactosidase (SA-β-gal) staining, revealing a markedly higher positivity rate in senescent cells (84.76 ± 3.15%) compared to proliferating controls (5.99 ± 1.27%), representing a 14.2-fold difference (t = 32.674, p < 0.001). Senescent cells exhibited characteristic morphological alterations, including enlarged cell bodies, increased diameter, and flattened phenotypes.(Figure 2A)

**Table 2.**
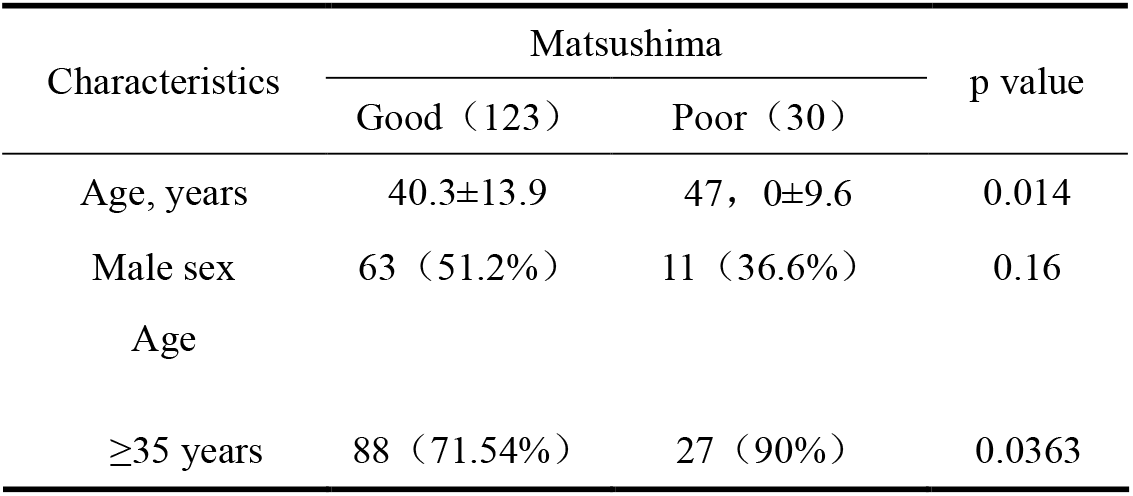
Association Analysis Between Postoperative Angiogenesis Quality and Baseline Characteristics. Notes: Age comparisons were analyzed using independent samples t-test; categorical variables were analyzed using Fisher’s exact test; p<0.05

| Characteristics | Matsushima |  | p value |
| --- | --- | --- | --- |
|  | Good (123) | Poor (30) |  |
| Age, years | 40.3±13.9 | 47, 0±9.6 | 0.014 |
| Male sex | 63 (51.2%) | 11 (36.6%) | 0.16 |
| Age |  |  |  |
| $\geq 35$ years | 88 (71.54%) | 27 (90%) | 0.0363 |

**Figure 1.**
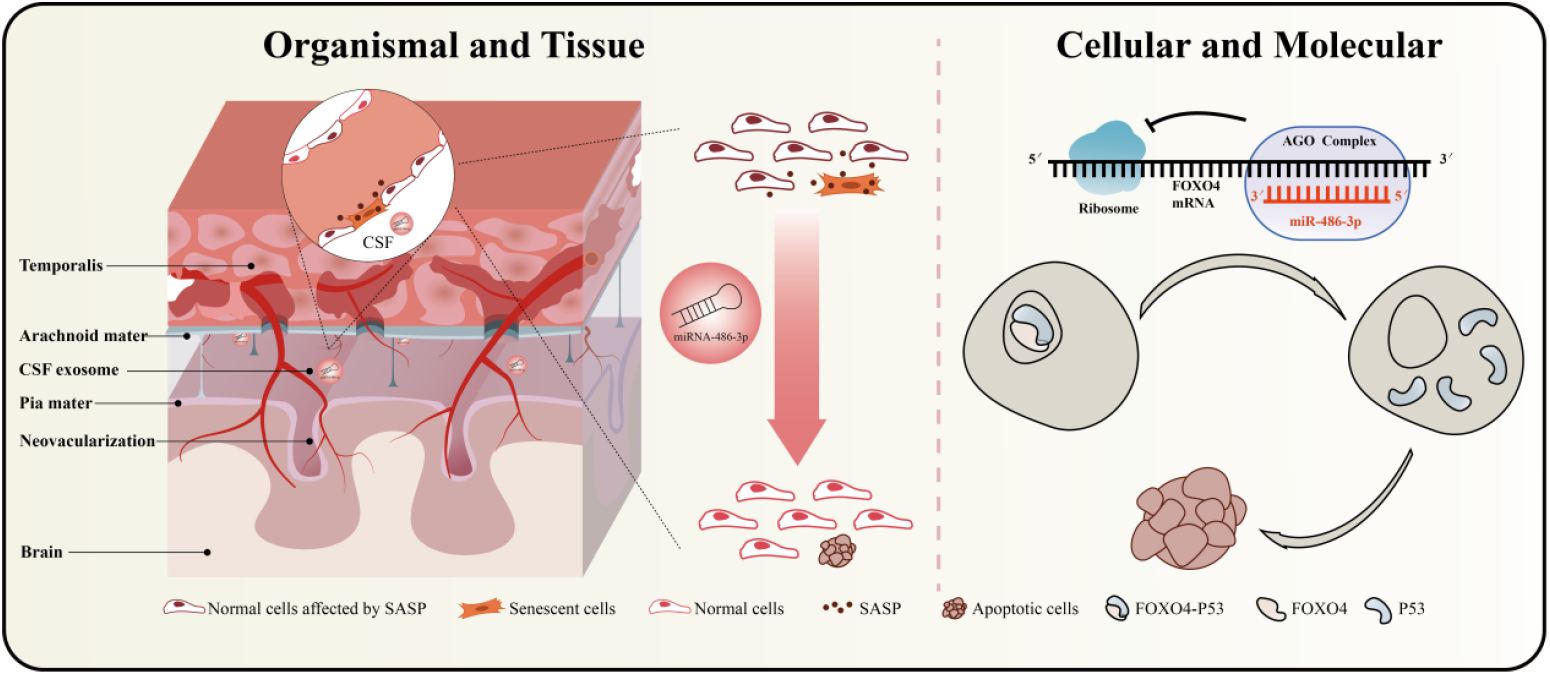
Research Hypothesis Diagram

**Figure 1.**
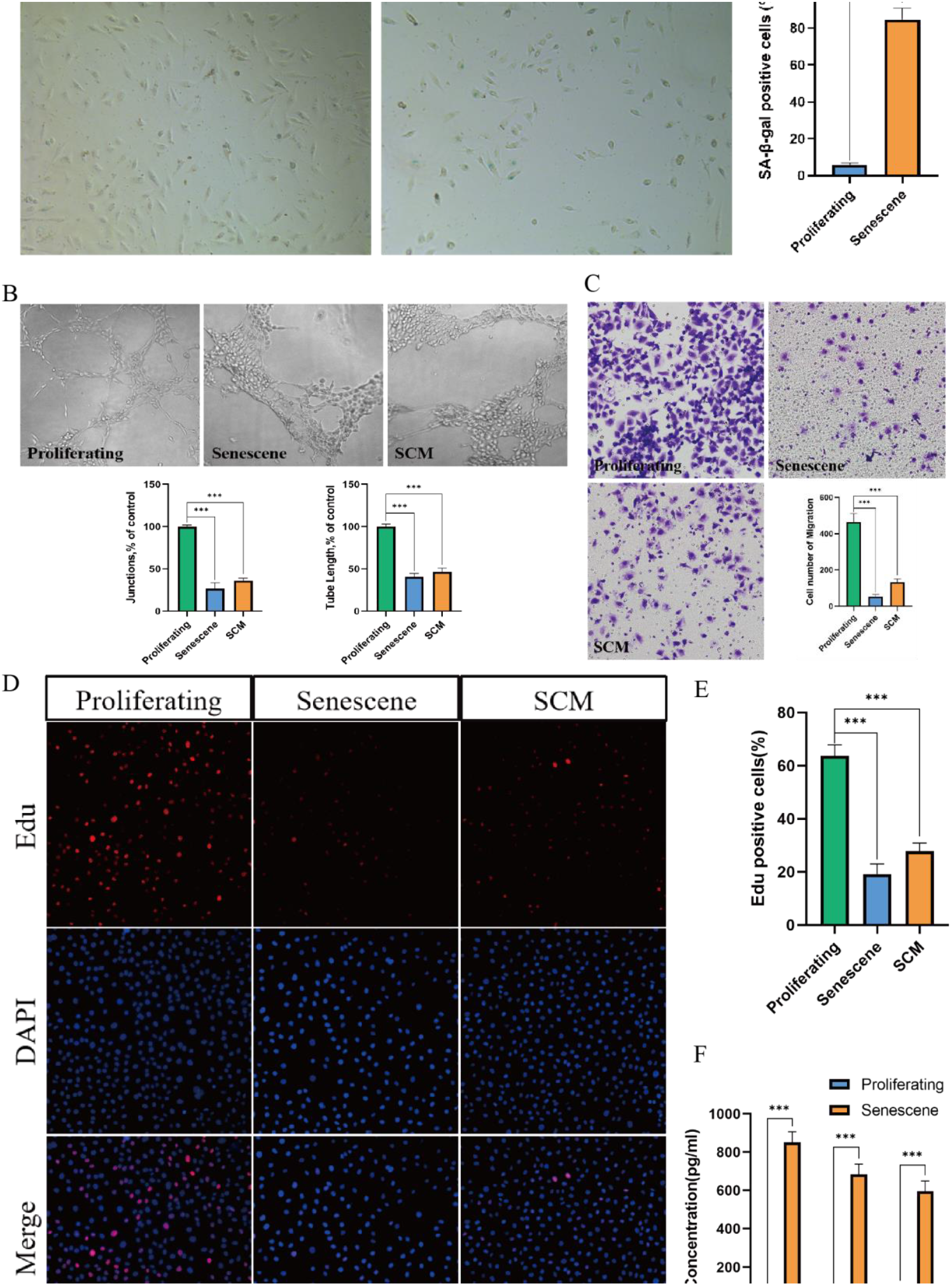
SASP-Mediated Angiogenesis Suppression by Senescent HUVECs

**Figure 2.**
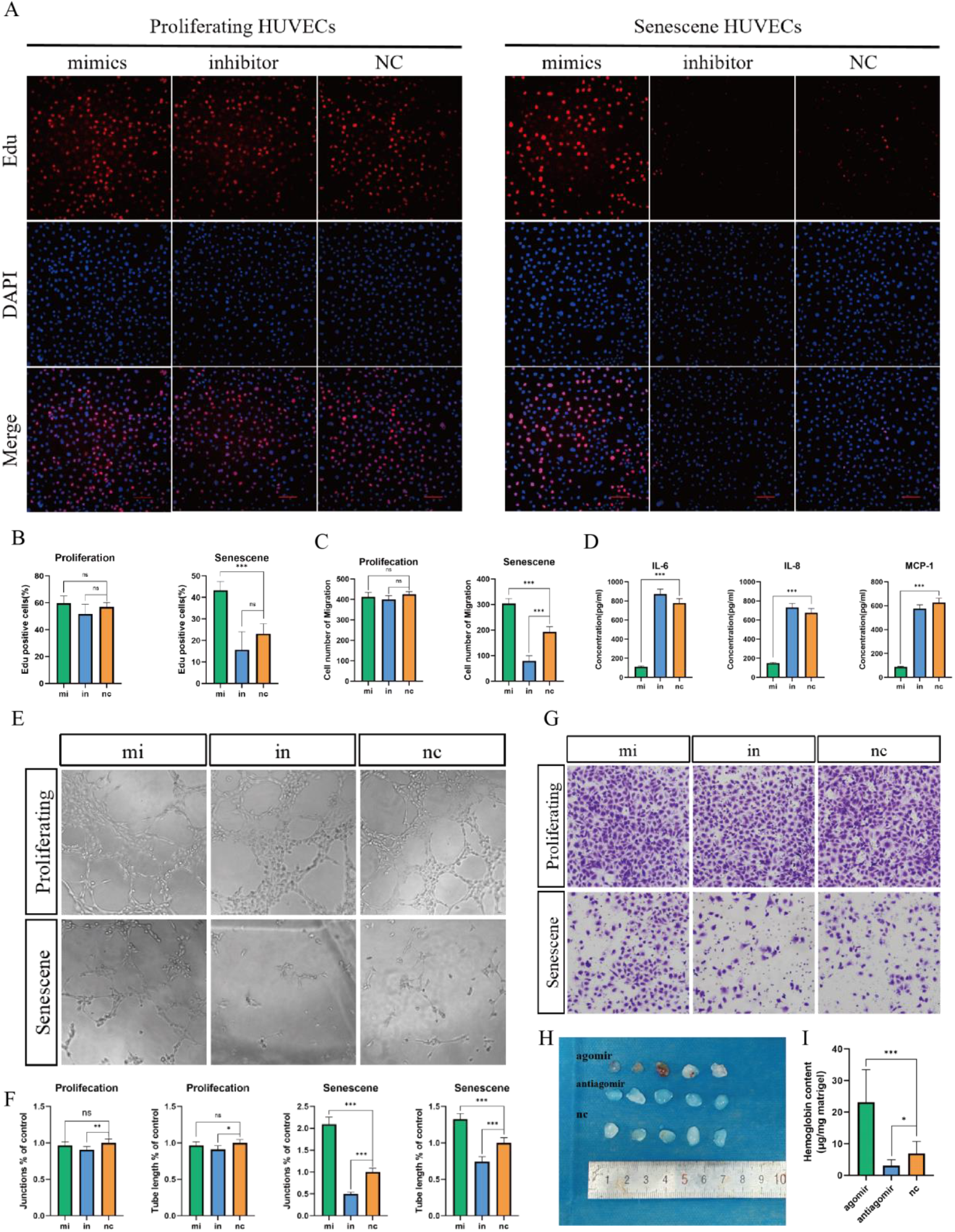
miR-486-3p Ameliorates Senescence-Associated Angiogenic Dysfunction

ELISA quantification demonstrated significantly elevated SASP factor secretion in senescent HUVECs (Figure 2F). All three SASP factors in senescent cells exceeded the 600 pg/mL threshold, confirming that serial passaging induces robust SASP secretion.

EdU assays revealed severe proliferation suppression in senescent cells (Figure 2D). Proliferating cells showed an EdU+ rate of 63.0 ± 4.1% (n = 6), which plummeted to 21.7 ± 4.8% in senescent cells (one-way ANOVA, p < 0.001). Proliferating cells treated with senescent conditioned medium (SCM) exhibited intermediate inhibition (28.3 ± 3.1%), significantly lower than controls (p = 0.0027) but higher than senescent cells (p = 0.032), indicating partial paracrine-mediated suppression(Figure 2E).

Transwell assays quantified migration deficits (Figure 2C). Proliferating cells displayed 464.83 ± 49.71 cells/field, whereas senescent cells showed an 88.6% reduction (53.17 ± 14.61 cells/field, p < 0.001). SCM-treated proliferating cells exhibited 132.83 ± 24.31 cells/field, a 71.4% decrease versus controls (p < 0.001) but higher than senescent cells (p = 0.0079), confirming soluble factor-mediated inhibition.

Matrigel tube formation assays demonstrated dual autocrine-paracrine inhibition of angiogenesis by senescence (Figure 2B). Senescent cells showed severe defects in vascular network complexity, while SCM-treated cells displayed intermediate impairment.

### 3.3 miR-486-3p Ameliorates Senescence-Associated Angiogenic Dysfunction

Based on our preliminary findings, cerebrospinal fluid levels of miR-486-3p, miR-92a-3p, miR-25-3p, and miR-155-5p differed significantly between revascularization-outcome groups in Moyamoya disease (MMD). While miR-92a-3p, miR-25-3p, and miR-155-5p have established roles in angiogenesis, miR-486-3p remains understudied, prompting its selection for functional analysis.

In proliferating HUVECs, miR-486-3p specifically impaired vascular network formation without affecting proliferation or migration:

In senescent HUVECs, miR-486-3p inhibition reduced proliferation (in: 15.72 ± 1.83% vs. NC: 21.34 ± 2.15%; p = 0.003), while mimics enhanced it (mi: 26.89 ± 3.04%; p = 0.019)(Figure 3A and 3B); Mimics increased migration by 52.6% (mi: 294.8 ± 21.3 vs. NC: 193.3 ± 21.5; p < 0.001), whereas inhibition reduced it by 58.7% (in: 79.8 ± 19.2; p < 0.001)(Figure 3C and 3G);Mimics doubled branch points (mi: 2.09 ± 0.17 vs. NC: 1.00 ± 0.09; +109.4%, p < 0.001) and increased total length by 32.5% (mi: 1.32 ± 0.08 vs. NC: 1.00 ± 0.08; p < 0.001). Inhibition halved branch points (in: 0.50 ± 0.05; -50.2%, p < 0.001) and reduced length by 25.8% (Figure 3E and 3F) ELISA revealed miR-486-3p-dependent regulation of SASP factors (IL-6, IL-8, MCP-1) in senescent HUVECs (Figure 3D).

**Figure 3.**
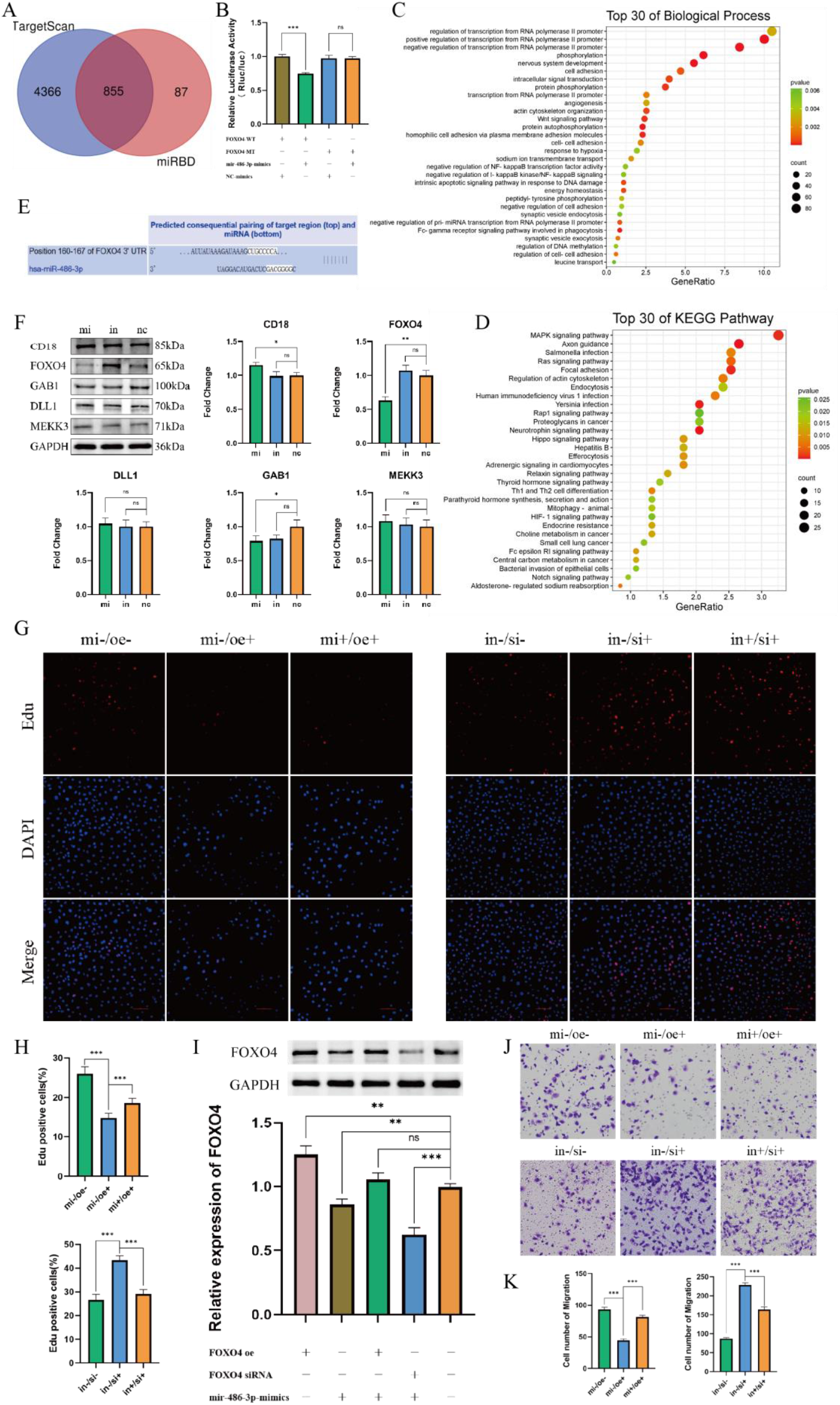
Mechanistic Elucidation of miR-486-3p in Regulating Senescent HUVEC Angiogenesis via FOXO4 Targeting Target Prediction & Pathway Analysis

Hydroxyurea-based assessment of vascular network maturity :Agomir group hemoglobin content increased 7.5-fold vs. antiagomir (p < 0.001) and 234.7% vs. NC (p = 0.002).Antiagomir reduced hemoglobin by 55.3% vs. NC (p = 0.041; one-way ANOVA: F = 18.94, p < 0.001)(Figure 3H)..

### 3.4 Mechanistic Elucidation of miR-486-3p in Regulating Senescent HUVEC Angiogenesis via FOXO4 Targeting Target Prediction & Pathway Analysis

Using downstream prediction websites TargetScan (http://www.targetscan.org/) and miRDB (http://mirdb.org/), we predicted the target genes of miR-486-3p. TargetScan and miRDB predicted 5263 and 942 downstream target genes, respectively. After performing a cross-analysis and taking the intersection of the two datasets, we obtained 855 common target genes, as shown in the figure below(Figure 4A).

Through Venn diagram analysis, we observed that 855 target genes were present in both websites’ predictions. Subsequently, we performed GO (Gene Ontology) and KEGG (Kyoto Encyclopedia of Genes and Genomes) enrichment analysis of these 855 genes using the DAVID website (https://david.ncifcrf.gov/). The top 30 biological processes and KEGG pathways were analyzed using RStudio (Version 4.0.2), as shown in the figure below(Figure 4C and 4D).

Based on the GO and KEGG results, we found that the target genes regulated by miR-486-3p may be involved in processes such as cell migration, cell adhesion, and neural system development, as well as regulating angiogenesis-related signaling pathways such as VEGFR signaling and Wnt signaling pathways. KEGG analysis also revealed that miR-486-3p participates in regulating Ras signaling, MAPK signaling, and Hippo signaling pathways, which are closely related to angiogenesis according to relevant literature. Therefore, we hypothesize that the target genes of miR-486-3p may participate in regulating the angiogenesis process. To further investigate the specific downstream target genes and pathways, we conducted subsequent studies.

In the GO analysis, we identified 29 angiogenesis-related genes. We then checked the number of binding sites and scores between these 29 genes and miR-486-3p using TargetScan, selecting the top five genes for Western Blot (WB) validation: ITGB2 (also known as CD18), GAB1, FOXO4, DLL1, and MAP3K3 (also known as MEKK3). Next, we transfected miR-486-3p mimics and miR-486-3p inhibitor into HUVECs using the same transfection method, and 48 hours later, extracted proteins for Western Blot experiments to validate the expression of the above genes and observe any changes in the expression of related proteins after interference, as shown in the figure below(Figure 4F).

The results showed that overexpression of miR-486-3p reduced FOXO4 expression, while inhibiting miR-486-3p expression led to a significant increase in FOXO4 expression. We therefore hypothesized that miR-486-3p can regulate FOXO4 expression. To investigate whether miR-486-3p exerts its effect by binding to FOXO4, we predicted potential binding sites between miR-486-3p and FOXO4 using the TargetScan website, as shown in the figure below(Figure 4E).

The results showed a binding site between miR-486-3p and FOXO4, leading us to hypothesize that miR-486-3p can regulate FOXO4 expression and exert its effects. To further verify the binding site, we performed a dual-luciferase reporter assay (Figure 4B).

Western blot analysis (Figure 4I) validated the targeted regulation of FOXO4 by miR-486-3p. FOXO4 overexpression significantly upregulated its protein levels (1.25 ± 0.05 vs. negative control [NC]: 1.00 ± 0.02; +25.0%, p < 0.01). In contrast, miR-486-3p mimics reduced FOXO4 expression (0.86 ± 0.04 vs. NC; −14.0%, p < 0.01). Co-transfection of miR-486-3p mimics with FOXO4 plasmid restored FOXO4 levels to 1.06 ± 0.04, showing no significant difference from NC (1.06 ± 0.04 vs. NC: p = 0.36) but a trend toward rescue compared to mimics alone (1.06 ± 0.04 vs. 0.86 ± 0.04; p = 0.067), confirming FOXO4 overexpression reverses miR-486-3p-mediated suppression.

EdU proliferation assays demonstrated miR-486-3p’s regulatory specificity(Figure 4G and 4H). In the FOXO4 overexpression model, miR-486-3p co-expression (mi+/oe+ group) partially reversed FOXO4-induced proliferation inhibition (EdU+ rate: 18.58 ± 1.25% vs. mi-/oe+ group: 14.87 ± 1.15%; p = 0.007). In the FOXO4 knockdown model, miR-486-3p inhibition (in+/si+ group) antagonized the pro-proliferative effect of FOXO4 knockdown (29.07 ± 1.72% vs. in-/si+ group: 43.32 ± 1.97%; p < 0.001), establishing miR-486-3p regulates proliferation homeostasis via FOXO4.

Transwell migration assays confirmed direct miR-486-3p-FOXO4 targeting(Figure 4J and 4K). In the FOXO4 overexpression model, miR-486-3p co-expression (mi+/oe+ group) partially rescued migration inhibition (81.17 ± 2.93 vs. mi-/oe+ group: 44.17 ± 2.56 cells/field; p < 0.001). In the knockdown model, miR-486-3p inhibition (in+/si+ group) attenuated FOXO4 knockdown-induced migration enhancement (164.83 ± 6.15 vs. in-/si+ group: 232.17 ± 5.15 cells/field; p < 0.001). One-way ANOVA confirmed significance (overexpression model: F = 356.2; knockdown model: F = 589.4; both p < 0.001).

## Discussion

A large number of researchers are dedicated to exploring the factors influencing angiogenesis after bypass surgery. Storey et al. found that the spontaneous formation of compensatory collateral circulation before surgery is an important predictor of good angiogenesis after bypass surgery. Compared to the control group, the postoperative Matsushima grading was significantly higher (1.8 vs 1.5, p < 0.003). In China, Zhao Yuanli et al. proposed that hemorrhagic moyamoya disease is a predictor of poor outcomes after bypass surgery, while the rich collateral vessels formed by internal carotid artery occlusion before surgery are closely related to the formation of good collateral vessels after surgery. Additionally, factors such as age at the time of surgery and preoperative ischemia are also considered as predictors of good postoperative outcomes. Unlike biological markers, the aforementioned factors are imaging or clinical markers. Although they can guide clinical decision-making to some extent, they do not provide a deep explanation of the physiological mechanisms.

Many clinical studies have also found that common cytokines closely related to angiogenesis, such as VEGF, FGF, HGF, PDGF, etc., are highly expressed in the dura mater, temporal muscle, or superficial temporal artery in patients with moyamoya disease. Endothelial progenitor cells (EPCs) are also significantly increased in the circulatory system of moyamoya disease patients. Hayashi et al. used bilateral carotid artery ligation in rats to simulate moyamoya disease and found that platelet-derived growth factor (PDGF) plays an important role in angiogenesis after temporal muscle bypass surgery. Marushima et al. found in rat models that myofibroblast-mediated VEGF/PDGF-BB could better induce angiogenesis and improve cortical perfusion. These studies provide some reference for the clinical practice and mechanism exploration of moyamoya disease bypass treatment. However, the direct mechanisms of angiogenesis after indirect revascularization surgery are still poorly understood and cannot explain the impact of age on angiogenesis.

What role does miR-486-3p play in the development of collateral circulation? To address this question, we first conducted a literature review. We found that miR-486-3p promotes cell apoptosis in various tumor diseases. Chou et al. found that increasing miR-486-3p expression could induce apoptosis in oral cancer cells. Similar findings have been reported in glioblastoma, hepatocellular carcinoma, thyroid cancer, and retinoblastoma. Ouyang et al. found that miR-486-3p promotes fluoride-induced osteoblast proliferation and activation by targeting the TGF-β1/Smad2/3 signaling pathway to transcriptionally regulate CyclinD1. Cui et al. found that miR-486-3p can regulate nucleus pulposus cell (NPC) proliferation and inhibit cell migration by inducing the expression of hyaluronan-binding protein CEMIP, and inhibiting miR-486-3p expression promotes NPC proliferation in intervertebral disc degeneration to achieve therapeutic effects for disc herniation.

The role of miR-486-3p in angiogenesis has not been reported. So, what role does miR-486-3p play in the angiogenesis process? To address this issue, we conducted preliminary research. Using human umbilical vein endothelial cells (HUVECs) as the model cells, we performed overexpression and inhibition treatments of miR-486-3p. The results showed no significant changes in the tube formation and migration abilities of HUVECs. In clinical practice, we noticed that the angiogenesis rate after temporal muscle bypass surgery in adult moyamoya disease patients is significantly lower than that in children, and further age group statistics revealed that as age increases, the angiogenesis rate gradually decreases. We reasonably hypothesized that endothelial aging due to age affects angiogenesis after temporal muscle bypass surgery.

In this study, we first verified through cell experiments that miR-486-3p promotes tube formation and migration in senescent HUVECs. Subsequently, we performed bioinformatics analysis to predict the potential target genes of miR-486-3p, and conducted qPCR, WB, and dual-luciferase reporter assays to validate these targets, ultimately identifying FOXO4 as a candidate target gene. We then used ELISA and WB to detect the expression of senescence-associated secretory proteins, and performed β-galactosidase staining and flow cytometry to detect cell apoptosis, confirming that miR-486-3p induces apoptosis in senescent HUVECs, with no effect on normal HUVECs. Finally, we constructed both overexpression and knockout models of FOXO4 in vitro and in vivo to explore the effects of the miR-486-3p/FOXO4 axis on angiogenesis regulation and its potential mechanisms.

## Conclusion

This study reveals that miR-486-3p ameliorates post-revascularization angiogenic impairment in Moyamoya disease (MMD) by targeting FOXO4 to clear senescent endothelial cells and inhibit senescence-associated secretory phenotype (SASP) secretion. These findings provide a novel therapeutic target for age-related vascular diseases and suggest miR-486-3p as a potential biomarker for predicting revascularization efficacy and a candidate therapeutic molecule.

## Author Contributions

Gang Wang and Wenfeng Feng designed the project. Junda Chen and Yunyu Wen performed the experiments and drafted the manuscript. Siyuan Chen, Tinghan Long, Fangzhou Chen and Zhibin Wang assisted with the experimental and data analysis. Wenfeng Feng and Gang Wang revised the manuscript critically. All authors contributed to the article and approved the submitted version.

## Research Funding

This work was supported by the Natural Science Foundation of Guangdong Province (2024A1515012964) and the Guangdong Medical Science and Technology Research Foundation (A2024541).

## Data Availability Statement

The authors confirm that the data supporting the findings of this study are available within the article and its supplementary materials.

---

**Supply Table:** Literature Review of Good Angiogenesis Rates After Indirect Revascularization.

| Year | The Author | Number of cases | Age of patients | Surgical methods | Methods of FOLLOW-UP | Good rate |
| --- | --- | --- | --- | --- | --- | --- |
| 1992 | Yamada et al.[27] | 27 | 1-20(8.3) | EDAS | Angiography(Mat sushima grade) | 76% |
| 1997 | Tenjin et al.[26] | 7 | 1-8(3.1) | EDAS | Angiography | 71% |
| 2012 | Ahn et al.[25] | 13 | 4-18(11.1) | EDAS;EDAS+EGPS; EDAS+EGS | Angiography(Mat sushima grade) | 82.4% |
| 2014 | Lin et al.[24] | 36 | 18.1-50.9(28.3) | Pial synangiosis | Angiography(Mat sushima grade) | 84.8% |
| 2016 | Liu et al.[23] | 30 | 4-16(12.6±3.1) | EDAS | Angiography(Mat sushima grade) | 80% |
| 2017 | Ogawa et al.[21] | 15 | 18-64(41.6) | EDAS | DSA(Matsushima grade) | 77.3% |
| 2017 | Blauwblomm et al.[22] | 64 | 4.4-18(7±4.3) | MBH | Angiography(Mat sushima grade) | 82.9% |
| 2019 | Wang et al.[20] | 65 | 20-54(37.1±8.1) | EDAS | Angiography(Mat sushima grade) | 74.3% |
| 2020 | Ge et al.[19] | 54 | 36±10.7 | EDAS;EDGS;MBH | DSA(Matsushima grade) | 42.2% |

